# Distinct physiological responses of *Coccolithus braarudii* life cycle phases to light intensity and nutrient availability

**DOI:** 10.1101/2022.02.17.480838

**Authors:** Gerald Langer, Vun Wen Jie, Dorothee Kottmeier, Serena Flori, Daniela Sturm, Joost de Vries, Glenn M. Harper, Colin Brownlee, Glen Wheeler

**Author notes:** equal contribution (Gerald Langer, Vun Wen Jie).

## Abstract

Coccolithophores feature a haplo-diplontic life cycle comprised of diploid cells producing heterococcoliths and haploid cells producing morphologically different holococcoliths. These life cycle phases of each species appear to have distinct spatial and temporal distributions in the oceans, with the heavily-calcified heterococcolithophores (HET) often more prevalent in winter and at greater depths, whilst the lightly-calcified holococcolithophores (HOL) are more abundant in summer and in shallower waters. The haplo-diplontic life cycle may therefore allow coccolithophores to expand their ecological niche, switching between life cycle phases to exploit conditions that are more favourable. However, coccolithophore life cycles remain poorly understood and fundamental information on the physiological differences between life cycle phases is required if we are to better understand the ecophysiology of coccolithophores. In this study, we have examined the physiology of HET and HOL phases of the coccolithophore *Coccolithus braarudii* in response to changes in light and nutrient availability. We found that the HOL phase was more tolerant to high light than the HET phase, which exhibited defects in calcification at high irradiances. The HET phase exhibited defects in coccolith formation under both nitrate (N) and phosphate (P) limitation, whilst no defects in calcification were detected in the HOL phase. The HOL phase grew to a higher cell density under P-limitation than N-limitation, whereas no difference was observed in the maximum cell density reached by the HET phase at these nutrient concentrations. HET cells grown under a light:dark cycle divided primarily in the dark and early part of the light phase, whereas HOL cells continued to divide throughout the 24 h period. The physiological differences may contribute to the distinct biogeographical distributions observed between life cycle phases, with the HOL phase potentially better adapted to high light, low nutrient regimes, such as those found in seasonally stratified surface waters.

**Highlights:**

- *Coccolithus braarudii* life cycle phases exhibit different physiological responses.
- The heavily-calcified heterococcolithophores (HET) life cycle phase is more sensitive to high light.
- The lightly-calcified holococcolithophores (HOL) life cycle phase may be better suited to growth under low phosphate availability.

## Introduction

Coccolithophores, unicellular marine algae, form a composite extracellular matrix called the coccosphere, comprised of organic material and intricately shaped calcite platelets, the coccoliths (Young *et al*., 2003). Coccolithophores are important primary producers (Poulton *et al*., 2007) and facilitate organic carbon export to the deep ocean through the ballasting effect of their coccoliths (Klaas & Archer, 2002; Ziveri *et al*., 2007). They are also, together with foraminifera, the greatest contributors to pelagic carbonate sediments (Broecker & Clark, 2009). Coccolithophores thrive in most marine environments from coastal waters to open oceanic areas and high salinity tropical seas such as the Gulf of Aqaba (Ziveri *et al*., 1995; Baumann *et al*., 2004; Tyrrell *et al*., 2008). The ability of coccolithophores to inhabit such diverse environments may be due in part to their specialised life cycle, although this aspect of coccolithophore biology so far remains poorly understood.

Coccolithophores exhibit a haplo-diplontic life cycle in which they can exist in alternate haploid or diploid life cycle phases that can both undergo mitotic divisions. Apart from their ploidy, the two life cycle stages are morphologically distinct. In many species, the haploid phase is lightly-calcified, producing holococcoliths that are comprised of simple rhombic crystals, whereas the diploid phase is heavily-calcified, producing heterococcoliths comprised of elaborately shaped crystals (Young *et al*., 2003). The holococcolithophore (HOL) phase is also motile in many species, whilst the heterococcolithophore (HET) phase is non-motile, although this distinction is not true for all species (Frada *et al*., 2018). The HOL and HET phases of each species have distinct spatial and temporal distributions in the environment (Supraha *et al*., 2016). Meta-analyses have revealed that HET tend to be most abundant in winter and at greater depths in the water column, whilst HOL are often relatively dominant in summer, particularly in surface waters (Malinverno *et al*., 2009; D’Amario *et al*., 2017). These distribution patterns strongly suggest that each life cycle phase occupies their own ecological niche, with the HOL phase most suited to nutrient-poor conditions with higher irradiances and temperatures, and the HET phase better suited to environments with lower irradiances and temperatures, and higher nutrient concentrations (de Vries *et al*., 2021). Whilst these distinct distribution patterns between life cycle phases can be observed in many species, it is not the case for all species. In several lineages, such as the Noelaerhabdaceae and Pleurochrysidaceae, the haploid phase is not calcified (Frada *et al*., 2018). As a consequence, our knowledge of the abundance and distribution of the haploid phase in these species remains very limited.

Improved knowledge of the differential physiology of HOL and HET life cycle phases is required to fully understand the ecology and biogeography of coccolithophores. The vast majority of laboratory studies on coccolithophore physiology have focused on the HET phase with only relatively few studies directly comparing both life cycle phases (von Dassow *et al*., 2009; Rokitta *et al*., 2011). These comparative studies are primarily in *Emiliania huxleyi*, which is the most abundant species in modern oceans. However, *E. huxleyi* is not an ideal species for the broader study of coccolithophore life cycles. The HOL phase is not calcified and very little is known about its environmental distribution (Frada *et al*., 2012). Moreover, genomic studies suggest that many *E. huxleyi* isolates exhibit life cycle modifications and may have lost the ability to revert to the HOL phase (von Dassow *et al*., 2015). The HET phase of *E. huxleyi* often forms large blooms in stratified surface waters, typified by high irradiances and low nutrient concentrations (Zondervan, 2007). This ability of *E. huxleyi* HET to thrive in nutrient-depleted surface waters sets it apart from many coccolithophore species, in which the HET phase is often most abundant in deeper waters with a higher nutrient concentration (D’Amario *et al*., 2017).

Further comparative studies of the physiology of HOL and HET phases are therefore required to understand their distribution patterns in the field. The heavily calcified HET phases of the closely related species *Coccolithus pelagicus* and *Coccolithus braarudii* play an important role in calcite production in the North Atlantic (Daniels *et al*., 2016). *C. pelagicus* HET is found in the Arctic and subarctic regions of the Atlantic and Pacific, whereas the larger *C. braarudii* is found primarily in coastal and upwelling systems in more temperate regions (Cachao & Moita, 2000; Saez *et al*., 2003; Daniels *et al*., 2016). A study of coccolithophore life cycle phases in the Mediterranean Sea noted that *C. braarudii* was primarily represented by its HOL phase, with the HET phase restricted to deeper waters in the South West Mediterranean (D’Amario *et al*., 2017). This suggests that *C. braarudii* HET is less suited to the warmer, oligotrophic regimes with a high irradiance that typify much of the Mediterranean. However, only very few studies have directly compared the physiology of *C. braarudii* life cycle phases. Houdan et al (2006) showed that a *C. braarudii* HOL isolate showed a less pronounced reduction in growth rate in low nutrient batch culture experiments (combined N and P limitation), suggesting that it is better suited to lower nutrient environments.

Whilst this is an intriguing observation, many further questions remain regarding the physiology of *C. braarudii* life cycle phases. Do they show differential sensitivity to light? How does growth under nutrient limitation compare when N or P are limited individually? Are there direct impacts of light and nutrient limitation on calcification and do these effects differ between the life cycle phases? The latter question is particularly relevant, as the degree of calcification is clearly a major physiological difference between HET and HOL life cycle phases. Coccolith morphology was unaffected by both N and P limitation in *C. leptoporus* HET cells (Langer *et al*., 2012), and by P limitation in *E. huxleyi* HET (Oviedo *et al*., 2014). However, *C. braarudii* HET cells showed an increase in coccolith malformations in response to P limitation (Gerecht *et al*., 2015). An increase in coccolith malformations was also observed when nitrate was replaced with urea as the N source (Benner, 2008). Moreover, it is unknown whether holococcolith crystal morphology is also influenced by nutrient limitation.

In this study, we have set out to compare the physiology of *C. braarudii* life cycle phases in response to changes in light and nutrient availability. We find that the *C. braarudii* HET exhibits a greater sensitivity to light, exhibiting reduced growth and defective coccolith morphology at high irradiances. We also find that the *C. braarudii* HOL is less detrimentally affected by nutrient limitation than the HET phase. These distinct responses may contribute to the environmental distribution of *C. braarudii* life cycle phases.

## Material and Methods

### Culture maintenance

Diploid (HET) *Coccolithus braarudii* (PLY182g – isolated from Western English Channel station E1) was obtained from the Plymouth Culture Collection, and the haploid (HOL) strain RCC3777 (isolated from the Bay of Biscay) from the Roscoff Culture Collection (http://roscoff-culture-collection.org/). Cells were grown in dilute batch cultures in a 15°C controlled temperature room in 0.2 µm filtered natural surface seawater sampled at E1 station off Plymouth, UK (50° 02.00’ N, 4° 22.00’ W) supplemented with modified K/2 media (http://www.roscoff-culture-collection.org/culture-media) with 220.5 µM nitrate and 9.05 µM phosphate. Incident light intensity was 20 µmol photons m^-2^ s^-1^ under a 16:8 h light:dark (LD) cycle. Both cultures were kept in exponential growth phase (500 to 30,000 cell ml^-1^) monitored by cell counts using Sedgewick Rafter chamber.

### Light experiment

Five light intensities (300, 50, 20, 10, 0 µmol photons m^-2^ s^-1^) were applied as determined by a LI-250A Photometer (LI-COR). Both life cycle stages were acclimated to 300, 50, 20 and 10 µmol photons m^-2^ s^-1^ respectively with an initial inoculum of 500 cells ml^-1^ for 5 generations. When cultures under 300, 50, 20, and 10 µmol photons m^-2^ s^-1^ light reached a cell density of 15,000 to 20,000 cell ml^-1^, the cells were inoculated into triplicate 40 ml culture flasks with an initial cell density of 500 cell ml^-1^. Cell density was determined daily by means of Sedgewick Rafter cell counting. Cells were harvested at 15,000 to 20,000 cells ml^-1^.

### Growth rate and discarded coccoliths to cell ratio

To avoid bias during sampling resulting from diel changes (Kottmeier *et al*., 2020), sampling was done four to eight hours after the start of light phase. The growth rates were calculated as described in (Kottmeier *et al*., 2020) by applying exponential linear regression analysis on the growth curves:

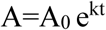

A and A_0_ are the final and initial cell densities; the growth rate is represented by k and t is time [days]; e = 2.71828183.

The discarded coccolith density was determined on harvest day for PLY182g only, because the holococcoliths produced by the haploid RCC3777 were too small for analysis. The number of discarded coccoliths per cell was determined by dividing the discarded coccolith density by the cell density.

### Cell Size Analysis

Cultures were harvested at 15,000 to 20,000 cell ml^-1^. For the diploid, 4 ml of culture was decalcified with 152 µl of 0.1 µM HCl for 2 minutes. The pH was later restored by the addition of 152 µl of 0.1 µM NaOH. Decalcified diploid cells were subsequently imaged using a mesolens microscope (McConnell *et al*., 2016). The haploid cells were also imaged with the mesolens without the need for decalcification as the protoplast can be clearly seen. For the protoplast diameter, 150 cells per biological replicate were measured using FIJI software.

### SEM microscopy of coccosphere and coccolith

Cultures were filtered onto Isopore membrane filters (diameter: 13mm; pore size: 0.8 µm) and rinsed with distilled water (1-2 seconds) to remove seawater. The filter was quickly dried with blue roll and left in a 60°C oven to dry overnight. The dried samples were mounted onto aluminium stubs and sputter coated with 10 nm Pt/Pd (Cressington, USA).

The number of visible coccoliths per coccosphere (n=21) was determined with a JEOL 6610 LV scanning electron microscope at a X4000 magnification. Coccolith morphology was classified into two categories, normal and malformed. We used a simplified categorisation compared to earlier studies (in this study ‘malformed’ contained the groups termed ‘minor’, ‘major’ and ‘type R’ (Langer *et al*., 2021)) as malformation levels were generally low and a high-resolution categorisation would not have provided additional information. A total of 200 coccoliths were analysed per sample. The visible coccoliths were taken to account for 75 % of the total coccolith quota (Hoffmann *et al*., 2015). Coccolith quota and coccolith length were used to calculate PIC quota for the HET phase. SEM imaging of intact cells for the lightly-calcified HOL phase is not possible as the holococcoliths and their coccospheres are usually disrupted during SEM sample preparation. PIC quotas therefore cannot be readily calculated for the HOL phase using this approach.

### Particulate organic and inorganic carbon (PIC & POC)

The measured protoplast diameter was converted into volume according to:

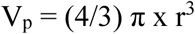

V_p_ is the protoplast volume and r is the radius of the cells. The calculated volume was converted into POC quota using the equation:

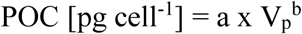

where a and b are constants (0.216 and 0.939 respectively) (Menden-Deuer & Lessard, 2000). To obtain the cellular PIC quota, the volume of the coccolith is required. The following equation was used.

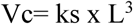

Here, Vc is the volume of coccoliths and can be estimated using coccolith length L and the shape constant ks (Young & Ziveri, 2000) which is 0.06 for *C. braarudii*. The cellular PIC quota is calculated from the following equation:

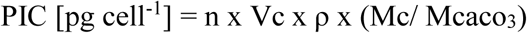

where n is the total number of coccoliths per cell including the discarded coccoliths; ρ is the calcite density of 2.7 pg µm^-3^ assuming coccoliths are pure calcite; Mc/ Mcaco_3_ is the molar mass ratio of C and CaCO_3_. Note that this calculation assumes a relatively constant coccolith shape and would not be appropriate for treatments that cause extensive coccolith malformations leading to a loss of the oval-shaped morphology. Although we did not specifically categorise the coccolith malformations observed in this study, malformations caused by high light were predominately minor malformations in which the oval-shape of the coccolith is maintained (see Fig. 2D). These would not be expected to greatly influence the calculation of cellular PIC quota.

### Dark Experiment

For the 0 µmol photons m^-2^ s^-1^ light treatment, an initial cell density of 15,000 cell ml^-1^ was inoculated into standard k/2 media in triplicate for both life cycle phases. Cell densities were measured daily for the initial 12 d to determine the survivability of the cells in the absence of light and then at 34 and 60 d.

### Nutrient Limitation Experiment

Both PLY182g and RCC3777 were grown in triplicate batch cultures in replete (control), P- and N-limiting medium at 15°C. The aim was to examine growth rate during exponential growth (a measure of growth at low nutrient concentrations prior to limitation), maximum cell density (a measure of the cellular quota of each nutrient) and the impacts on calcification. Cultures were grown in modified K/2 medium with 9.05 µM phosphate and 220.5 µM nitrate (control), initial phosphate of 1.5 µM (P-limiting) or an initial nitrate of 20 µM (N-limiting). The cultures were grown at an incident light intensity of 50 µmol photons m^-2^ s^-1^ on a 16:8 h light: dark (LD) cycle. Cell density was determined daily using a Sedgewick Rafter Cell. In the nutrient limited cultures, cells were harvested in the late exponential phase, which was determined in a pre-experiment. For the control cultures, cells were harvested in exponential phase at a similar cell density to avoid differential cell density effects. For growth rate determination, only the exponential part of the growth curve was included (2-5 d).

The response of cells to nutrient replenishment was also tested. Cells were grown under initially low N or P as described above. Upon reaching stationary phase (day 8), the cultures were diluted with the initial medium in a 1:1 ratio. Growth rates were determined in the subsequent growth phase after one day (two-point).

### Light microscopy

P and N- limited cells were imaged using a TMS inverted microscope with a 40x objective (Nikon, Japan), captured with an EOS 1200D camera (Canon, Japan), as well as a DMi8 inverted microscope with a 63x objective oil immersion objective (Leica, Wetzlar, Germany) and captured with an ORCA-Flash 4.0 camera (Hamamatsu, Japan).

### Photo-physiology measurements

Measurements of the photosynthetic efficiency of photosystem II (Fv/Fm) were made on dark-adapted (30 min) cells using an AquaPen-C device (Photon Systems Instruments, Drásov, Czechia). All measurements were made 5 d after inoculation in the experimental conditions, during the phase of exponential growth.

## Results

### Cellular responses to light

*C. braarudii* strains RCC3777 (HOL) and PLY182g (HET) were grown at light intensities ranging from 10 to 300 µmol photons m^-2^ s^-1^. While both life cycle phases showed light saturated growth at 50 µmol photons m^-2^ s^-1^, the specific growth rate of the HET phase significantly decreased at 300 µmol photons m^-2^ s^-1^ whereas the growth of the HOL phase remained close to maximum (Fig. 1A). The HET cells also displayed a reduction in estimated POC and PIC production at 300 µmol photons m^-2^ s^-1^ relative to 50 µmol photons m^-2^ s^-1^ (Fig. 1B-C). In contrast, estimated POC production in HOL cells continued to increase with light intensity up to 300 µmol photons m^-2^ s^-1^ (Fig. 1B). These results suggest that the HOL phase is better adapted to high light than the HET phase in terms of growth and POC production rates.

**Figure 1:**
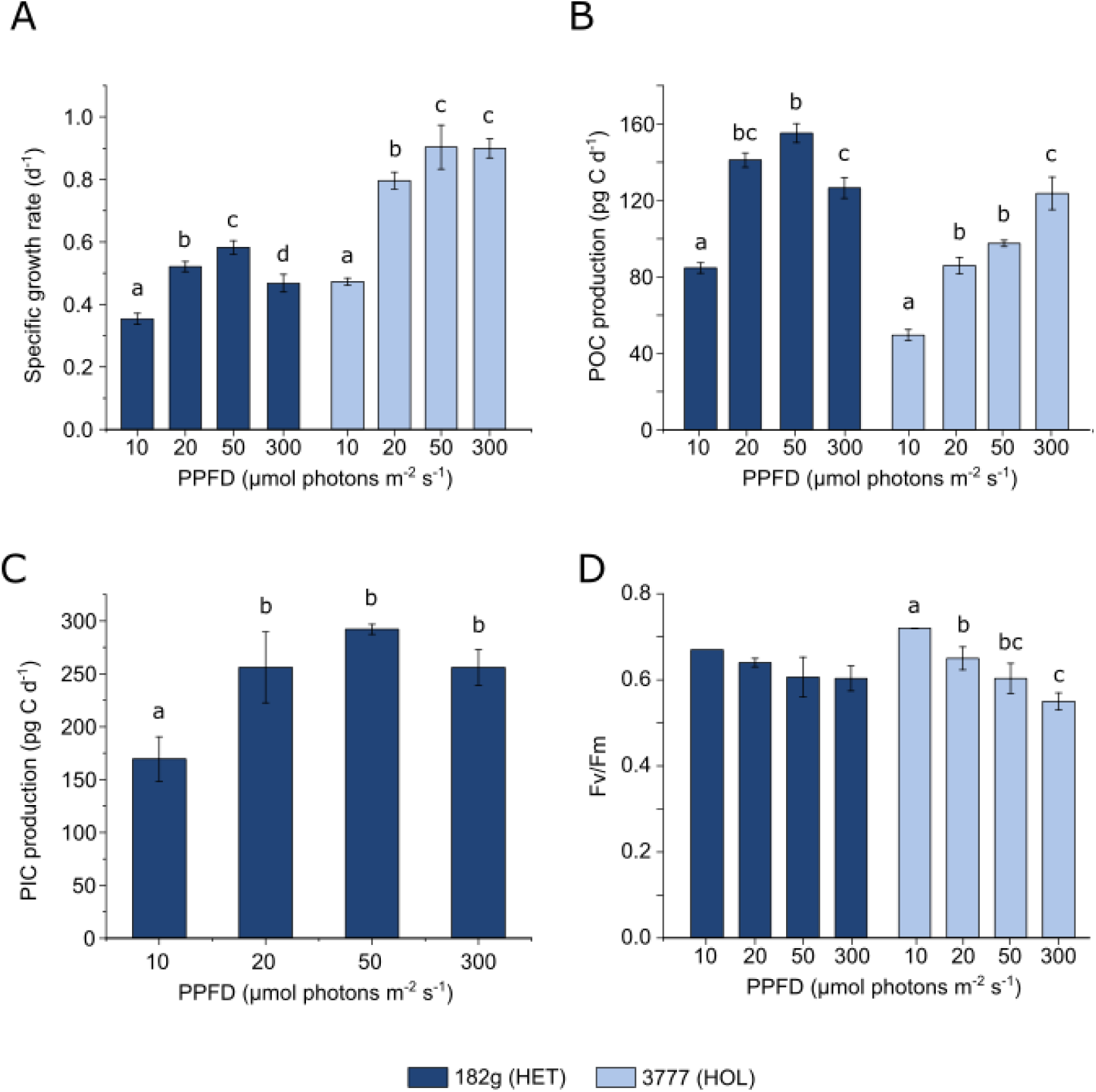
Physiological responses of *C. braarudii* life cycle phases to changes in irradiance. **A)** Growth of heterococcolithophore (HET) PLY182g and holococcolithophore (HOL) RCC3777 at different irradiances. **B)** Estimated particulate organic carbon (POC) production of HET and HOL life cycle phases. **C)** Estimated particulate inorganic carbon (PIC) production of the HET phase. Please note that there are no PIC data for the HOL phase (see Material and Methods for details). **D)** Maximum quantum efficiency of PSII (Fv/Fm). In all cases n=3, error bars = SD, letters represent treatments that are statistically different within each life cycle phase (one way ANOVA, Holm-Sidak post hoc test).

To examine whether these physiological responses were the consequence of photoinhibitory damage to the light harvesting mechanisms, we determined the photosynthetic efficiency of photosystem II in the dark-adapted state (Fv/Fm). HOL cells displayed a trend of decreasing Fv/Fm with increasing light (Fig. 1D). HET cells also exhibited a small decrease in Fv/Fm at higher irradiances, although this was not statistically significant. This suggests that the impaired growth at 300 µmol photons m^-2^ s^-1^ in the HET phase is not a consequence of substantial photoinhibition of PSII.

To further characterize the impact of high light on calcification in the HET phase, we first measured the number of discarded coccoliths in each treatment. The morphology of the heterococcoliths is important for the structural integrity of the coccosphere, as they overlap with neighbouring coccoliths. Malformed coccoliths often fail to integrate into the coccosphere, so the number of discarded coccoliths is an indication of the extent of coccolith malformations. Cells grown at 300 µmol photons m^-2^ s^-1^ showed a clear increase in the number of discarded coccoliths relative to all other light levels (Fig. 2A). We then analysed heterococcolith morphology in more detail using scanning electron microscopy (SEM). HET cells grown at 300 µmol photons m^-2^ s^-1^ also showed a significant increase in the proportion of coccolith malformations compared to all other light levels (Fig. 2B-D). Holococcoliths are substantially more fragile than heterococcoliths and they tend to disintegrate during preparation for SEM analysis. Therefore, we were only able to analyse the morphology of the crystals, as opposed to the morphology of the coccolith as a whole or the integrity of the coccosphere. Crystal morphology in holococcoliths from HOL cells grown at 300 µmol photons m^-2^ s^-1^ was not different from the crystals grown at the lower light intensities (97.8 ± 1.7% crystals exhibited normal morphology, compared to >96% in all other samples) (Supplementary Fig. 1). Overall, these results suggest that calcification in the HET phase is sensitive to high light, leading to a decrease in PIC production and impacts on coccolith morphogenesis and the structural integrity of the coccosphere.

**Figure 2:**
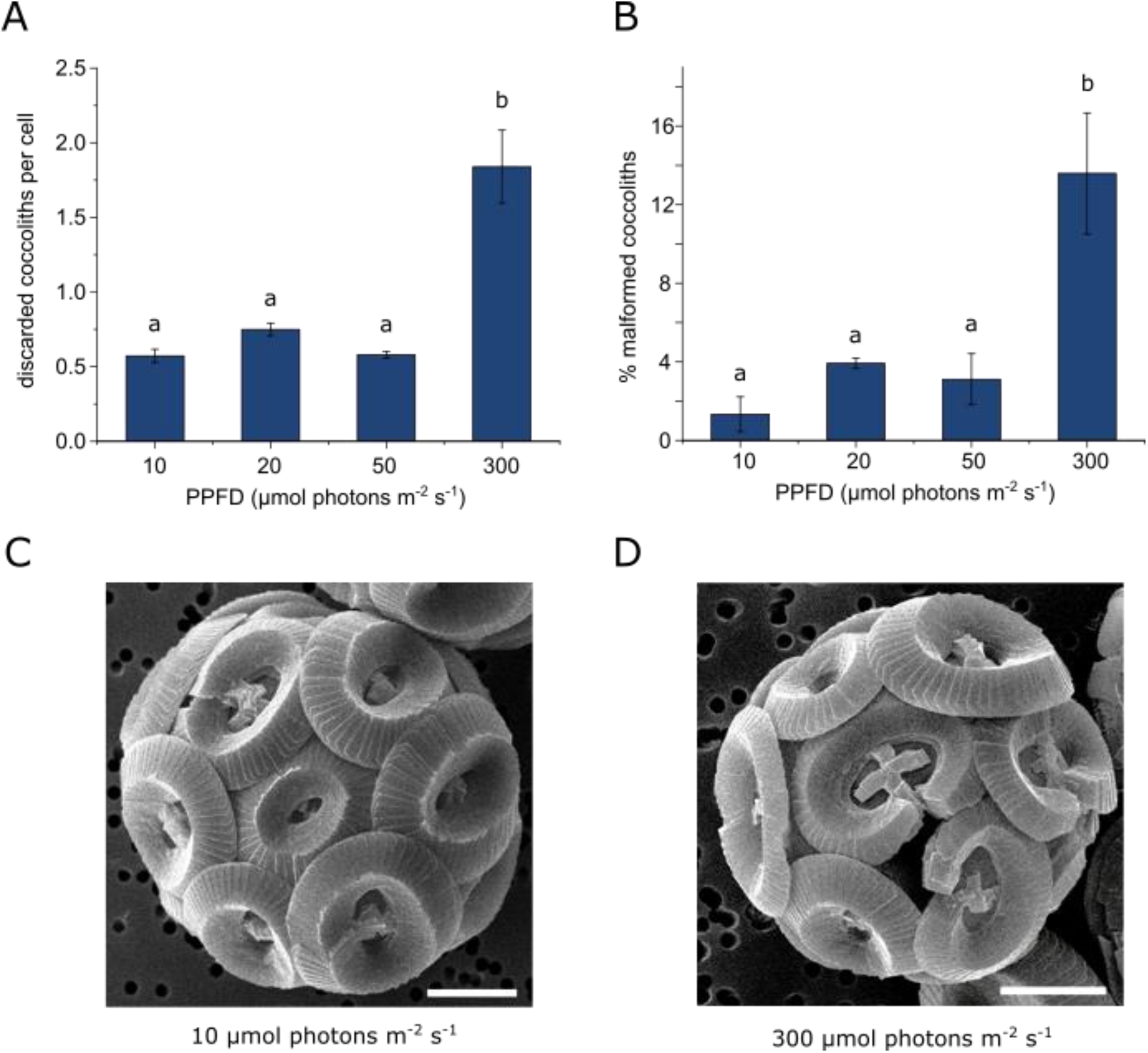
Effect of irradiance on calcification in *C. braarudii* HET. **A)** Number of discarded heterococcoliths per 182 g HET cell during growth at different irradiances, determined by light microscopy. The discarded coccoliths that do not integrate correctly into the coccosphere are often malformed. n=3, error bars = SD, letters represent treatments that are statistically different (one way ANOVA, Holm-Sidak post hoc test). **B)** Effect of light on coccolith morphology. The percentage of malformed heterococcoliths was determined by SEM analysis. **C)** Representative SEM image of a 182g HET cell grown at 10 μmol photons m^-2^ s^-1^. Bar = 2 μm. **D)** Representative SEM image of a 182g HET cell grown at 300 μmol photons m^-2^ s^-1^. Note the presence of malformed coccoliths. Bar = 2 μm.

The HOL phase may therefore be better suited to higher light environments than the HET phase. This is consistent with the distribution of *C. braarudii* life cycle phases in the field, both spatially (depth) and temporally (seasonal irradiance) (D’Amario *et al*., 2017). However, it is notable the HET phase did not grow better than the HOL under very low light conditions (10 µmol photons m^-2^ s^-1^). We therefore examined the ability of both life phases cycle stages to survive in an extended period of darkness, which may be beneficial for cells occupying a deeper position in the water column. We found that cell density decreased rapidly in the HOL phase after 7 days in darkness, with cell density approaching zero after 10 days (Fig. 3). In contrast, the cell density of the HET phase did not decrease for the entire 60 day duration of the experiment (Fig. 3). Microscopic observation indicated that HET cells remained pigmented throughout. Thus, the HET phase is able to survive in the absence of a photosynthetic supply of energy during extended periods of darkness.

**Figure 3:**
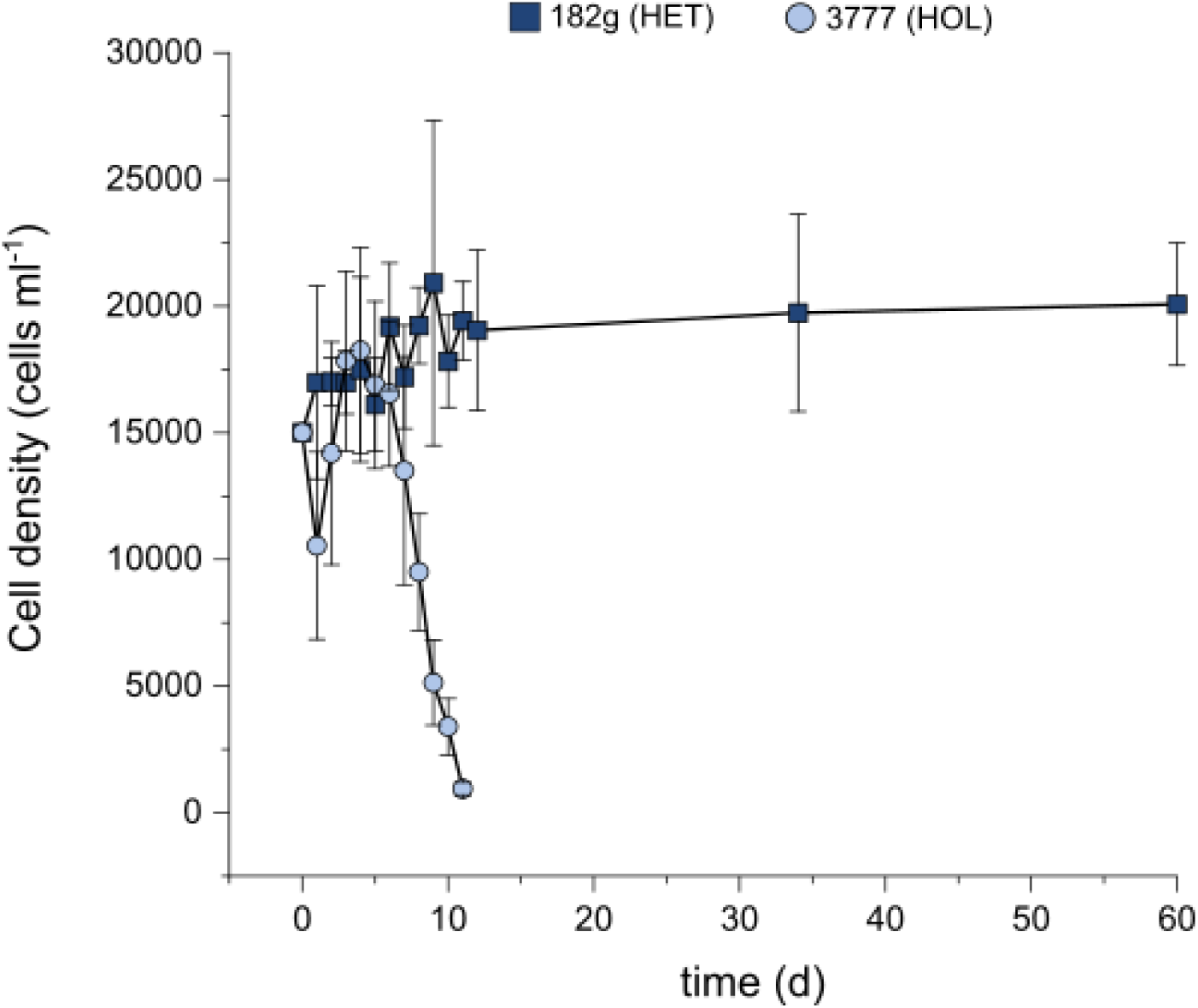
Survival of *C. braarudii* in darkness. Cell density of 182g HET and 3777 HOL life cycle phases during a prolonged dark period. Cells were grown to mid exponential phase and then placed in continuous darkness at T=0 for the duration of the experiment. n=3, error bars = SD.

### Response to nutrient limitation

We next examined the growth of the HET and HOL phases of *C. braarudii* under nutrient limiting conditions. We grew *C. braarudii* life cycle phases in batch or semi-continuous culture with limiting concentrations of N or P (20 μM or 1.5 μM respectively). These culturing approaches have distinct advantages and disadvantages when studying nutrient limitation (Langer *et al*., 2012; Langer *et al*., 2013; Gerecht *et al*., 2015). In batch culture experiments, nutrient concentrations decline throughout the experiment, as they are taken up by the cells and not replenished in the media (Langer *et al*., 2013). However, the maximum cell density reached can be used as an indication of the minimum cell quota of a limiting nutrient (Perrin *et al*., 2016). The HOL phase grew to a higher cell density under P limitation relative to N limitation, whereas the cell densities obtained by the HET phase were very similar under both N- and P-limitation (Fig. 4A-B).

**Figure 4:**
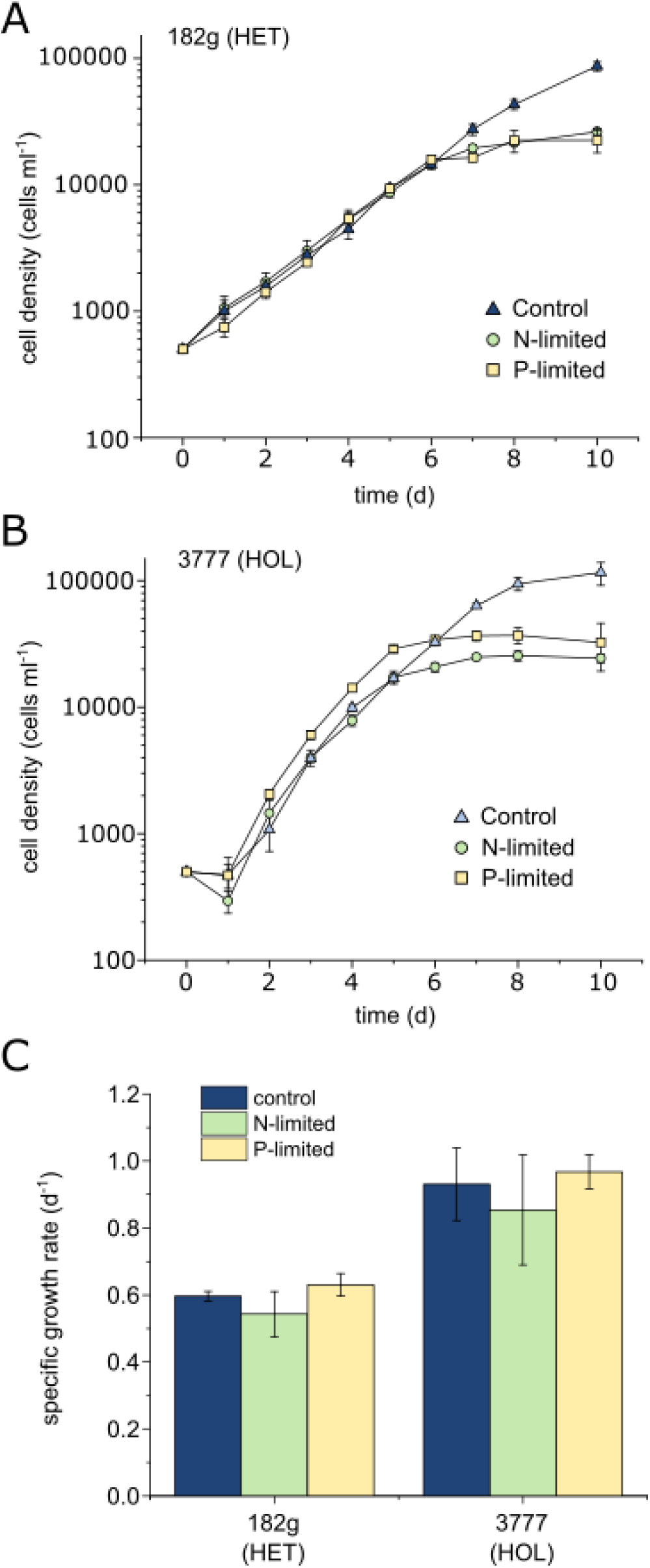
Response of *C. braarudii* life cycle phases to nutrient limitation. **A)** Growth of 182g HET in nutrient limited batch cultures. N-limited initial nitrate concentration 20 µM, P-limited initial phosphate concentration 1.5 µM. n=3, error bars = SD. **B)** Growth of 3777 HOL in nutrient limited batch cultures. **C)** Specific growth rates calculated from the period of exponential growth prior to nutrient limited growth (2-5 d for all, except 3777 HOL P-limited and N-limited, 2-4 d). No significant differences were found between treatments in each life cycle phase (one way ANOVA).

Growth rate is another important ecological parameter, although this can only be determined reliably in the exponential phase of a batch culture. This puts considerable restrictions on the interpretation, because the nutrient limitation effects are comparatively weak in the exponential phase, with the cells only becoming severely nutrient depleted in the transition phase or early stationary phase (Langer *et al*., 2012). Growth rate in the exponential phase of a batch culture therefore reflects the ability of cells to grow in low but not necessarily limiting nutrient concentrations. There was no significant effect of low N or P on the growth rate of HOL or HET cells (Fig. 4C).

We next examined growth in response to nutrient replenishment. Batch cultures that were exhibiting N- or P-limited growth (day 8) were diluted with an equal volume of fresh media containing low concentrations of N or P (20 μM or 1.5 μM respectively). We found that the growth rate of HET cells was restored to similar values following either N or P replenishment (74.7% or 82.3% respectively of the nutrient replete control), whereas in HOL cells, replenishment of P led to a much greater increase in growth rate (128.6% of a nutrient replete control) compared to N replenishment (97.0% of control) (Fig. 5). This suggests that the HOL phase is able to respond more quickly to a restoration of nutrient availability following nutrient limitation, particularly when P is resupplied. These are likely to be beneficial attributes in areas of low or episodic nutrient supply.

**Figure 5:**
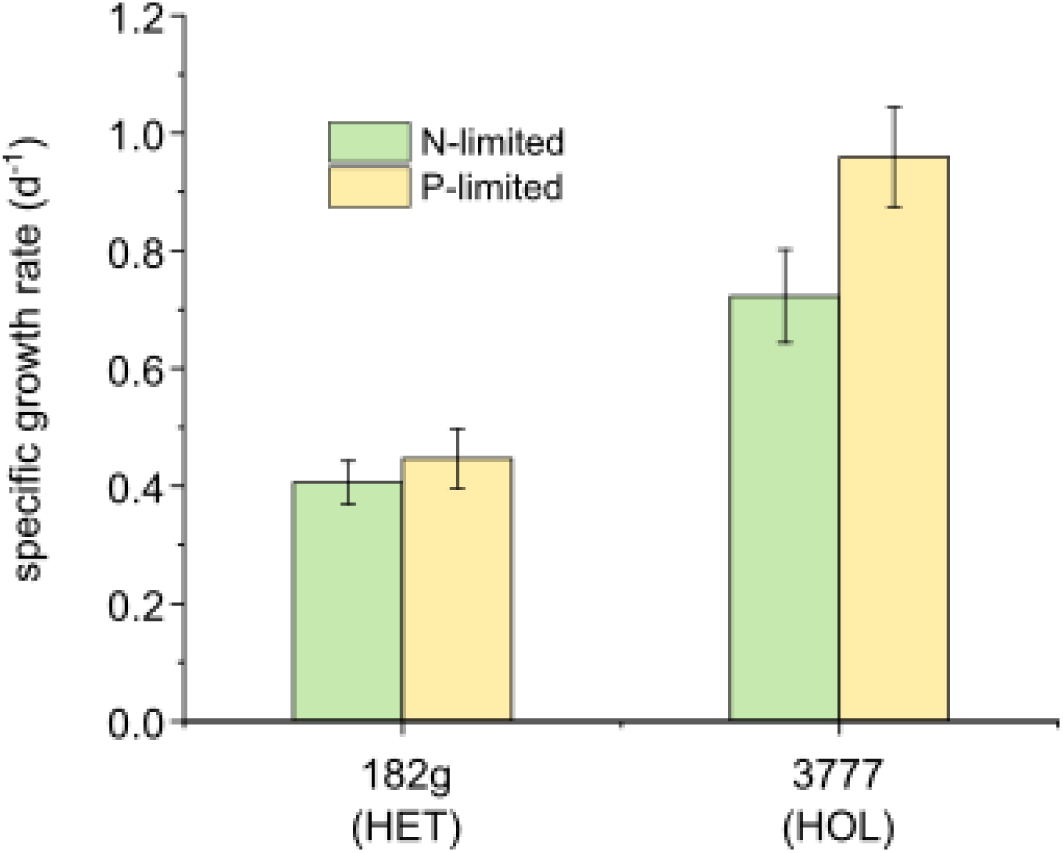
Growth rate of *C. braarudii* life cycle phases following nutrient replenishment. 182g HET and 3777 HOL life cycle phases were grown in N-limited (20 μM) or P-limited (1.5 μM) batch cultures (as shown in Figure 4). At day 8 when growth rate had slowed, cultures were diluted with an equal volume of fresh media containing 20 µM N (for N-limited cultures) or 1.5 µM P (for P-limited cultures). Growth rate was determined 24 h after nutrient replenishment. n=3, error bars = SD.

The impact of nutrient limitation on heterococcolith morphology has been a long standing question. Analysis of coccoliths in environmental samples led to suggestions that nutrient limitation may result in coccolith malformations (Okada & Honjo, 1975; Kleijne, 1990). Experimental studies subsequently confirmed this hypothesis for P limitation in the *C. braarudii* HET (Gerecht *et al*., 2015). We therefore analysed heterococcolith morphology in nutrient-limited cells from our batch culture experiments. We found that P- as well as N-limitation caused an increase in the percentage of discarded coccoliths per cell and a substantial increase in the percentage of coccoliths exhibiting malformations (Fig. 6A-B). P limitation had a more severe effect than N limitation for both of these metrics. Whilst we could not directly compare these parameters in the HOL phase, we examined the crystal morphology of holococcoliths. All treatments displayed predominately normal crystals (Fig. 6C, Supplementary Fig. 2). However, it was noticeable that HOL cells grown under P limitation had more complete coccospheres than control or N-limited cells (Fig. 6D). Although we were not able to quantify this effect directly, it again points to the tendency of the HOL phase to perform better under P limitation than N limitation.

**Figure 6:**
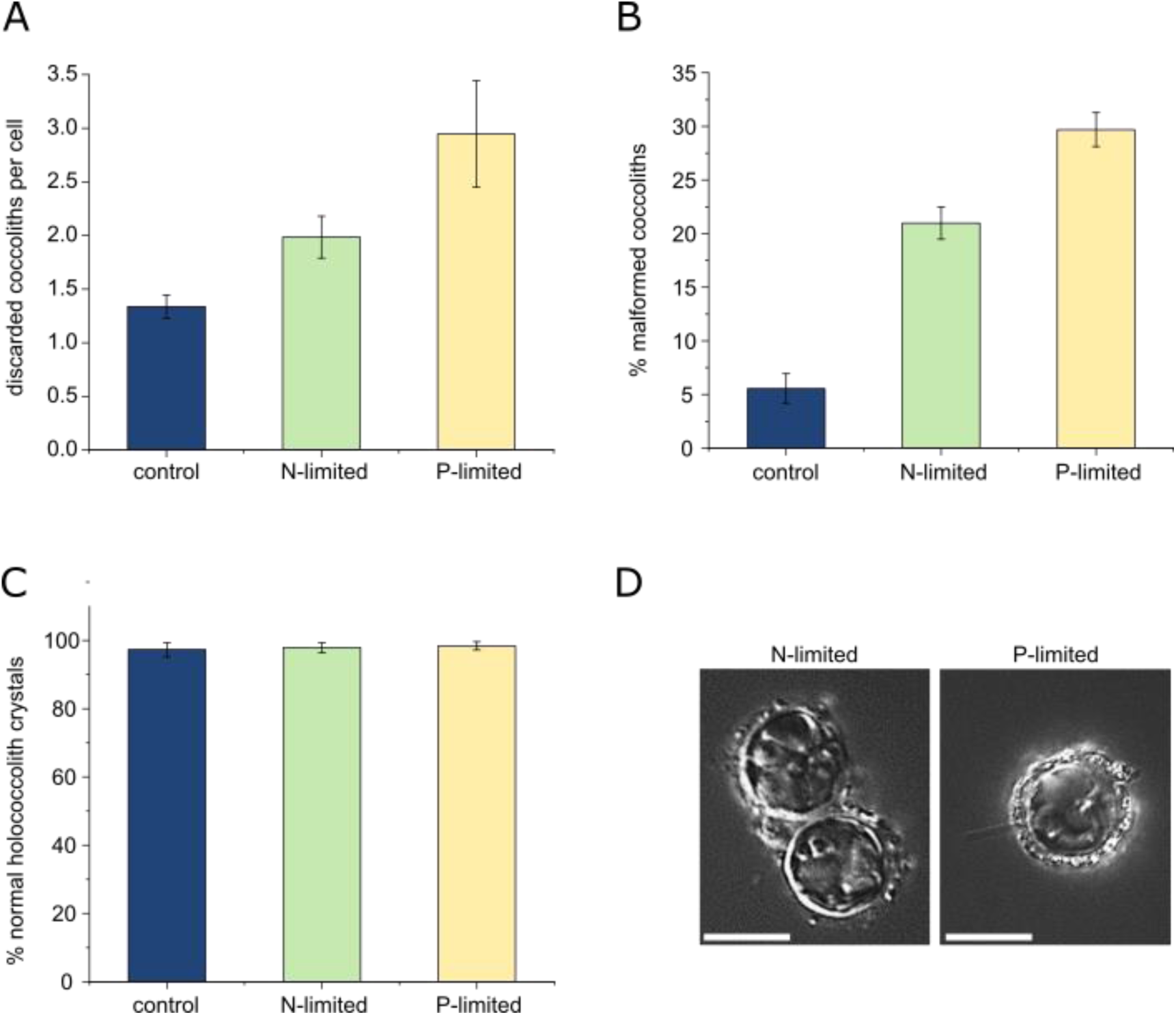
Effect of nutrient limitation on calcification in both life cycle phases. **A)** Number of discarded heterococcoliths per 182 g HET cell during nutrient limited growth, determined by light microscopy. n=3, error bars = SD. Coccoliths that fail to integrate into the coccosphere are often malformed. **B)** Effect of nutrient limitation on 182g HET coccolith morphology. The percentage of malformed heterococcoliths was determined by SEM analysis. **C)** Effect of nutrient limitation on the morphology of crystals within holococcoliths of 3777 HOL. Crystal morphology was examined rather than the holococcolith itself, as the integrity of holococcoliths is not preserved during preparation for SEM analysis. **D)** Light micrographs of N- and P-limited 3777 HOL *C. braarudii* showing a larger, more complete coccosphere in P-limited cells, compared to N-limited cells. Scale bars = 10 μm.

### Timing of cell division in the diel cycle

*E. huxleyi* HET cells grown under a light-dark cycle become synchronised, with cell division occurring primarily in the dark phase although it can continue for several hours after the onset of the light phase (Muller *et al*., 2008; Kottmeier *et al*., 2020). Calcification occurs primarily in the light, with the rate of calcification tightly correlated to the percentage of cells in the G1 phase of the cell cycle (Muller *et al*., 2008). We therefore examined the nature of cell division in HOL and HET phases of *C. braarudii* to see if this was influenced by the light-dark cycle (16:8). Similar to previous observations of *E. huxleyi* HET (Kottmeier *et al*., 2020), *C. braarudii* HET showed only a small increase in cell density during the light phase, with the majority of cell division occurring in the dark phase (Fig. 7). In contrast, HOL cells showed a linear increase in cell density throughout the light-dark cycle. Continuous, rather than synchronised, cell division in the HOL phase could also explain why the HOL phase is able to show an improved growth rate under repeated nutrient replenishment in the semi-continuous culture (Fig. 5).

**Figure 7:**
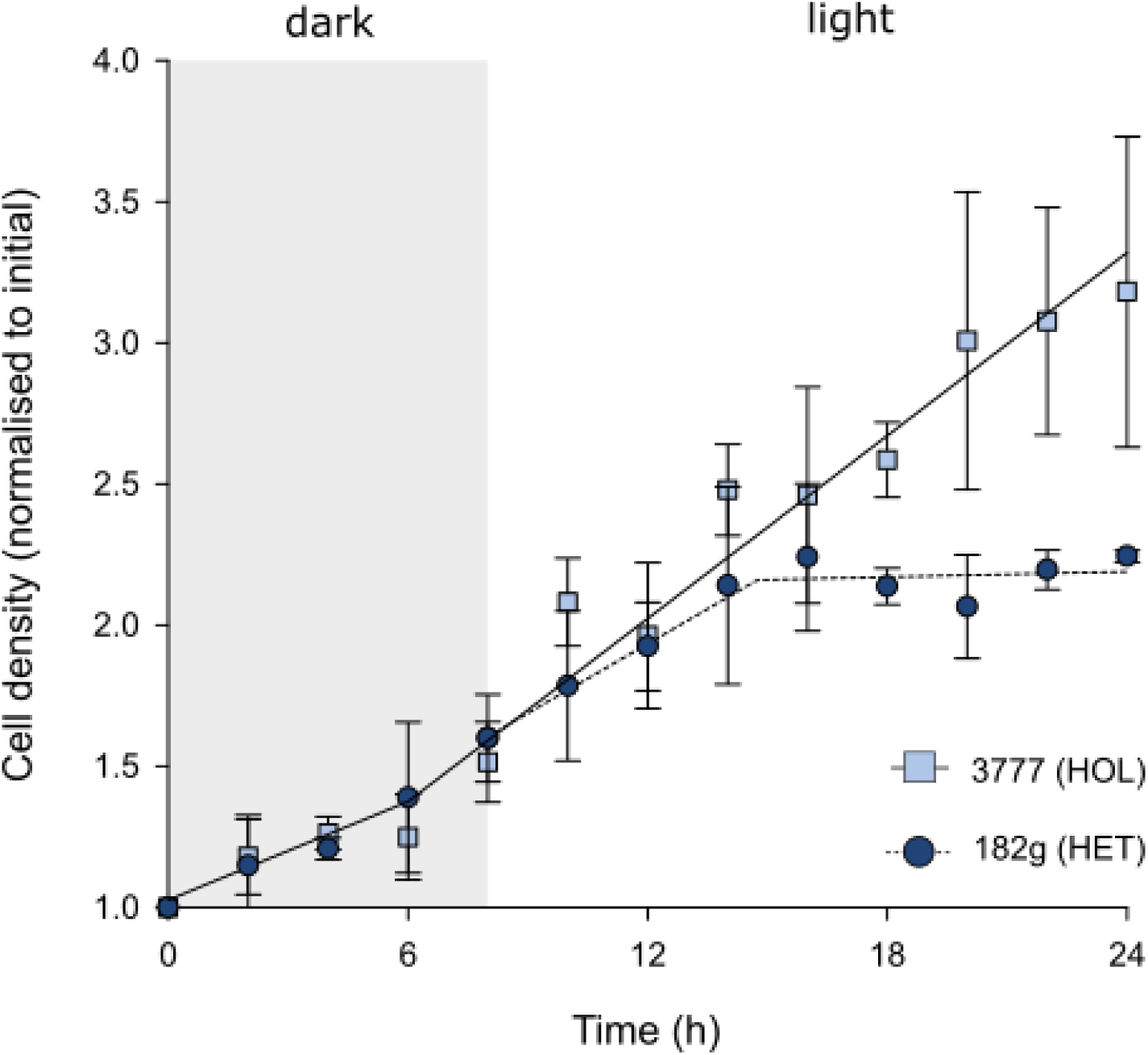
Timing of cell division throughout the diel cycle. Relative cell density of 182g HET and 3777 HOL cultures are shown throughout the diel cycle (16:8, light:dark). The cell densities are shown normalised to cell density at T=0 to demonstrate the relative changes in growth rate. The HET culture divides primarily in the dark phase, with cell increasing only in the initial part of the light phase. In contrast, the cell density of the HOL phase increases steadily throughout the diel cycle. A piecewise two segment linear regression was performed to illustrate changing trends across the 24 h period between the life cycle phases. n=3, error bars=SD.

## Discussion

Our results demonstrate that *C. braarudii* life cycle phases exhibit distinct responses to changes in irradiance. The HET phase was notably sensitive to higher irradiances, showing an optimum of 50 μmol photons m^-2^ s^-1^ across the tested range. This suggests that the natural populations of *C. braarudii* HET could encounter irradiances above optimal levels in surface waters in the N. Atlantic, although they would be much less likely to encounter excess light lower in the water column as PAR attenuates rapidly with depth (Poulton *et al*., 2010). The greater sensitivity of HET phase to light is therefore in keeping with its expected distribution in the water column (D’Amario *et al*., 2017). As the HET phase did not show a decline in the performance of photosystem II under high light, the direct cause of the greater sensitivity of the HET phase to light remains unclear. One possibility is that the high demand for dissolved inorganic carbon for photosynthesis may compete with the carbon supply for calcification (Bach *et al*., 2013; Bolton & Stoll, 2013).

The response of other coccolithophore species to light has been studied most extensively in *E. huxleyi. E. huxleyi* HET blooms primarily in surface waters during mid-summer and high irradiance has been identified as an important contributory factor in bloom formation (Nanninga & Tyrrell, 1996; Tyrrell & Merico, 2004). Accordingly, laboratory studies indicate that *E. huxleyi* (HET) is highly tolerant to higher irradiances (Ragni *et al*., 2008; Loebl *et al*., 2010; Gafar & Schulz, 2018). The *E. huxleyi* HOL phase is more sensitive to light, exhibiting photoinhibition at light intensities above 400 µmol photons m^-2^ s^-1^ (Houdan *et al*., 2005). The greater sensitivity of *E. huxleyi* HOL to light is therefore in sharp contrast with the findings from *C. braarudii* and seem to run counter to the expectation from many coccolithophore species that HOL prefer higher light compared to the HET. However, our very limited knowledge of the ecological distribution of the *E. huxleyi* HOL means that we cannot determine whether the physiological differences between life cycle phases in this species reflect adaptation to different environments. *Gephyrocapsa oceanica* is closely related to *E. huxelyi* and its HET phase is similarly tolerant of high irradiances (Zhang *et al*., 2015). There are relatively few laboratory studies of light tolerance in the coccolithophore species found at greater depths, although *Scyphosphaera apsteinii* HET appears to be low-light adapted with growth optima around 100 µmol photons m^-2^ s^-1^ (Drescher *et al*., 2012; Gafar *et al*., 2019).

The *C. braarudii* HET phase showed a remarkable ability to survive in darkness. This ability could be ecologically relevant in life cycle phases that occupy a deeper position in the water column or inhabit high latitude regions, which experience extended periods of darkness in winter months. Whilst *C. braarudii* HET is not typically found in very high latitudes, the closely related *C. pelagicus* HET is abundant in Artic and subarctic waters. We do not yet know why *C. braarudii* HET cells are able to survive longer in darkness. In extended darkness, one strategy polar phytoplankton use to provide organic nutrients is mixotrophy (McKie-Krisberg & Sanders, 2014; Stoecker & Lavrentyev, 2018). However, previous studies suggest that bacterivory is most likely to be exhibited by the *C. braarudii* HOL phase rather than the HET phase, although this ability has also been observed in the HET phase of some other placolith bearing coccolithophores (Houdan *et al*., 2006; Avrahami & Frada, 2020). It therefore appears unlikely that dark survival of the HET is due to improved ability to acquire nutrients via bacterivory relative to the HOL phase. Dark survival of the HET phase may therefore be due to another aspect of physiology, such as an improved capacity to utilise stored lipids, carbohydrates or proteins as organic carbon sources, as observed in some polar diatoms (Schaub *et al*., 2017). A recent study demonstrated that dark survival in HET coccolithophores was supported by osmotrophy of organic carbon compounds (Godrijan *et al*., 2022). The trends observed in the physiological responses of the *C. braarudii* life cycle phases to light are therefore broadly consistent with their different seasonal and geographical distributions.

There were also indications from our experiments that the HOL phase was better suited to low nutrient environments, although this was observed primarily under P-limitation rather than N-limitation. In particular, the higher cell density achieved by HOL cells under P-limitation suggest a low minimum cell quota for P that may be advantageous in environments where P availability is low. Transcriptomic and metabolomic studies of N-limitation in *E. huxleyi* indicate an overall decrease in growth and biosynthetic capacity in both life cycle phases, presumably due to the lack of available N for protein synthesis (Rokitta *et al*., 2014; Wordenweber *et al*., 2018). Whilst P-limited *E. huxleyi* cells displayed some shared metabolic responses with N-limited cells (Rokitta *et al*., 2016), it is thought that a primary consequence of P-limitation is the inability to synthesise P-rich nucleic acids, which limits growth but not necessarily other aspects of metabolism (Wordenweber *et al*., 2018). Some of the major consequences of P-limitation in *E. huxleyi* HET are lipid remodelling to reduce phospholipid content and the activation of P-scavenging enzymes such as alkaline phosphatase (Shemi *et al*., 2016). It is interesting that significant differences in the expression of these P-acquisition genes was observed between *E. huxleyi* life cycle phases, which may contribute to the greater ability of HOL cells to resist P-limitation (Rokitta *et al*., 2016). The lower DNA content in the haploid genome of HOL cells may also contribute to a lower P-requirement that would provide an advantage under P-limitation (Rokitta *et al*., 2016). A combination of these factors may therefore contribute to the improved growth of the *C. braarudii* HOL under P limitation observed in our experiments.

A notable consequence of both light and nutrient stress in the HET phase was the disruption of coccolith morphology. Both N and P limitation had a detrimental effect on heterococcolith crystal morphology while they had no effect on holococcolith crystal morphology. The response to light is surprising because there is no obvious mechanistic connection between the morphogenetic machinery and light intensity. The impaired morphogenesis in the HET *C. braarudii* under 300 µmol photons m^-2^ s^-1^ is ecologically relevant because light intensities in the surface mixed layer can be higher than 300 µmol photons m^-2^ s^-1^ (Nanninga & Tyrrell, 1996), and aberrant coccolith morphology might impair coccosphere architecture and cell division (Walker *et al*., 2018). The findings suggest that holococcolith formation does not require the cellular mechanisms within heterococcolith formation that are sensitive to excess light or nutrient limitation. It can be hypothesized that the HOL crystal shaping machinery is simpler and is less easily disturbed by differing demands on cellular resources such as inorganic carbon, N and P. This hypothesis fits the observation that holococcolith crystals are all rhombic and of the same relatively small size (species independent), while heterococcolith crystals display complex shapes and sizes that can even vary within a single coccolith (Young *et al*., 2003). The relative insensitivity of holococcolith morphogenesis to environmental stress might be one advantage of the HOL phase in certain habitats.

Calcification is clearly a major physiological difference between the *C. braarudii* life cycle phases and it may also contribute to the observed differences in the timing of cell division. Both heterococcoliths and holococcoliths are primarily produced in the light phase of the diel cycle (Paasche, 1969). Production of each single large heterococcolith in the HET phase takes approximately three hours, whereas holococcolith production is much more rapid (Taylor *et al*., 2007; Langer *et al*., 2021). It seems likely that the extensive rearrangement of the cytoskeleton required for cell division would interfere with its essential role in heterococcolith formation (Langer *et al*., 2010; Durak *et al*., 2017) and intracellular heterococcoliths are not observed in dividing HET cells (Walker *et al*., 2018). Therefore heterococcolith formation and cell division are temporally separated in the HET phase, with the former occurring primarily in the light and the latter restricted to the dark phase of the diel cycle. This distinction is not found in the HOL phase, which appears to divide throughout the diel cycle. The ability to generate and secrete mature holococcoliths very rapidly means that cell division would not cause a lengthy disruption of the calcification process. It is possible that the lack of synchronisation of cell division in the HOL phase may allow for enhanced growth rates under certain conditions, such as the response to pulsed nutrient additions in low nutrient regimes.

Our results identify some important physiological differences between two *C. braarudii* life cycle phases, 182g (HET) and RCC3777 (HOL). Our results broadly support the conclusion of Houdan *et al* (2006) who also observed lower sensitivity of the HOL phase to macro-nutrient limitation. The physiological differences are striking because they are closely aligned to expectations based on the differing environmental distributions of *C. braarudii* life cycle phases (Cros & Estrada, 2013; Supraha *et al*., 2016; D’Amario *et al*., 2017). By exhibiting distinct physiology between life cycle phases, *C. braarudii* and other coccolithophores can occupy a broader ecological niche (de Vries *et al*., 2021). However, it is important to note that our findings are based on the observation of two distinct environmental isolates. Whilst both strains were isolated from a similar geographic location (NE Atlantic), it is possible that they have a degree of inherent variability in their sensitivity to various stressors. This is largely unavoidable in laboratory comparisons of phytoplankton strains, but this is an important caveat to our findings. Future studies examining further isolates or ideally isogenic HET-HOL pairs isolated from the same culture (Houdan *et al*., 2006) will be needed to further verify our observations.

## Acknowledgements

We thank Malcolm Woodward (Plymouth Marine Laboratory, UK) for help with nutrient analyses.

## Author contributions

G. Langer: original concept, all experimental analyses, drafting and editing manuscript; VW Jie: all experimental analyses; D Kottmeier: cell division experiment; S Flori: cell division experiment; D Sturm: analysis of nutrient limitation experiments; J de Vries: analysis of nutrient limitation experiments; G Harper: electron microscopy; C Brownlee: drafting and editing manuscript: G Wheeler original concept, drafting and editing manuscript.

